# Linear Z-line-like alignment of capping protein in obliquely striated muscle of the nematode *C. elegans* suggests that dense bodies are not equivalent to Z-lines

**DOI:** 10.1101/2025.11.24.690283

**Authors:** Shoichiro Ono, Emily Nickoloff-Bybel, Kennosuke Kurimaru, Kanako Ono

## Abstract

Many invertebrates have obliquely striated muscles, in which neighboring thin and thick filaments are staggered and aligned in an oblique manner. This type of muscle allows force production over a wide range of lengths and is beneficial for soft-bodied animals. Unlike cross-striated muscles of vertebrates, most of obliquely striated muscles lack distinct Z-lines and, instead, have dense bodies. Because the dense bodies are located in the middle of the I-bands and contain α-actinin, the dogma is that dense bodies are equivalent to the Z-lines anchoring the actin barbed ends. However, we present evidence that the barbed ends of sarcomeric actin filaments in the nematode *Caenorhabditis elegans* body wall muscle are aligned in a linear Z-line-like arrangement without converging at the dense bodies. Colocalization of F-actin and ATN-1/α-actinin was minimal. Furthermore, CAP-1, an α-subunit of capping protein/CapZ, was linearly aligned in the middle of the I-bands without concentration at the dense bodies. This linear CAP-1 alignment was maintained in the absence of ATN-1. These results demonstrate that the actin barbed ends are not directly anchored to the dense bodies. Depletion of the capping protein subunit, CAP-1 or CAP-2, caused embryonic or larval lethality with severe actin disorganization in the body wall muscle, indicating that barbed-end regulation by capping protein is essential for sarcomere assembly. These results contradict the current view of the sarcomere organization in *C. elegans* muscle and suggest a new model of a linear Z-line-like arrangement of actin barbed ends.

**Significance Statement:**

- Without clear evidence, there has been a notion that actin filaments are directly anchored to the dense bodies in *C. elegans* striated muscle.
- Capping protein localizes in a linear Z-line-like alignment in *C. elegans* muscle without concentrating at the dense bodies, indicating that the actin barbed ends are not directly anchored at the dense bodies.
- Depletion of capping protein causes severe sarcomere defects in embryos and larvae indicating a critical role of capping protein in sarcomere assembly.

## Introduction

In striated muscle, actin, myosin, and other regulatory proteins are arranged in a sarcomeric manner and adapted to specialized function to generate contractile forces (Sanger *et al*., 2017; Prill and Dawson, 2020). Vertebrates and some invertebrates have transversely cross-striated muscles, in which the Z-discs (or lines) are aligned perpendicularly to the filament axis and anchor the ends of the thin filaments (Burgoyne *et al*., 2015; Wang *et al*., 2021). In the Z-discs, α-actinin cross-links actin filaments (Ebashi and Ebashi, 1965; Maruyama and Ebashi, 1965; Masaki *et al*., 1967), and capping protein (CP) [also known as β-actinin (Maruyama, 1965) or CapZ (Casella *et al*., 1986)] caps and stabilizes the barbed ends of actin filaments. Fish α-actinin and CP directly interact *in vitro* and are proposed to act together as a part of an anchoring complex for the thin filaments (Papa *et al*., 1999).

However, many invertebrates, in at least 18 different phyla, have obliquely striated muscles, in which neighboring thin and thick filaments are slightly staggered and aligned in an oblique manner (Toida *et al*., 1975; Taylor-Burt *et al*., 2018; Taylor-Burt *et al*., 2025). The oblique angle to the axis of the thin and thick filaments, termed stagger angle, changes during muscle contraction and relaxation: the stagger angle is 2∼6 ⁰ at a relaxed state but increases to ∼30 ⁰ at a contracted state (Hanson and Lowy, 1957; Rosenbluth, 1967; Knapp and Mill, 1971). This change occurs due to the process called “shearing”, in which neighboring thin and thick filaments slide during contraction, such that the degree of stagger is decreased when muscle is contracted (Rosenbluth, 1967; Knapp and Mill, 1971). Combination of actomyosin sliding and shearing in obliquely striated muscles allows shape changes over much wider range of lengths than simple actomyosin sliding in cross-striated muscles (Taylor-Burt *et al*., 2025). Therefore, these physiological characteristics of obliquely striated muscles are considered beneficial for soft-bodied animals.

The body wall muscle of the nematode *Caenorhabditis elegans* is obliquely striated muscle and has been used as a model to study structure and function of contractile apparatuses (Benian and Epstein, 2011; Ono, 2014). This muscle lacks distinct Z-lines but instead has dense bodies, which are finger-like projections of ∼0.2 µm in diameter extending ∼1.0 µm from the plasma membrane into the cytoplasm (Francis and Waterston, 1985). Because the dense bodies are located in the middle of the I-bands and enriched with ATN-1/α-actinin, there has been a notion that the dense bodies are analogous to the Z-lines where the ends of sarcomeric actin filaments are directly inserted and anchored (Moerman and Fire, 1997; Cox and Hardin, 2004; Lecroisey *et al*., 2007; Gieseler *et al*., 2017). However, to date, there has been no clear demonstration of the direct anchorage of the actin-filament ends to the dense bodies. Early ultrastructural studies on the body wall muscle of a large nematode species, *Ascaris lumbricoides*, showed that small clumps of thin filaments, termed “Z bundles”, are distinct from the dense bodies and distributed in a band-like pattern in the middle of the I-bands (Rosenbluth, 1965), but such structures have not been confirmed in *C. elegans* muscle. In this study, we examined a spatial relationship between ATN-1/α-actinin and CAP-1, an α subunit of CP, as markers for the dense bodies and actin barbed ends, respectively, and found that CP-marked actin barbed ends are minimally localized to the dense bodies but, rather, distributed in a linear Z-line-like pattern in the middle of the I-bands. Thus, in contradiction to the previous notion, we propose that the sarcomeric actin filaments are linked indirectly to the dense bodies in *C. elegans* obliquely striated muscle.

## Results and Discussion

### Actin is not concentrated at or near the dense bodies in *C. elegans* body wall muscle

In the *C. elegans* body wall muscle, the dense bodies are ∼0.2 µm in diameter and spaced with gaps of 1.0 ∼ 1.5 µm (Francis and Waterston, 1985). If the dense bodies are the sites where all the barbed ends of sarcomeric actin filaments are directly anchored, a single dense body needs to accommodate all the ends of actin filaments spread over 1.0 ∼ 1.5 µm within the I-band. If this is the case, there should be substantial accumulations of actin at or near the dense bodies. However, we found no such accumulations of F-actin at or near the dense bodies, when localization patterns of F-actin and ATN-1/α-actinin, as a marker for the dense bodies, were compared (Fig. 1A, B). The patterns of sarcomeric actin in the *C. elegans* body wall muscle appear different depending on the contraction status (Butkevich *et al*., 2015). When the muscle was relaxed, F-actin appeared in a chain-like pattern in which clear holes were present in the middle of the I-bands (Fig. 1A). These clear holes corresponded to the dense bodies as indicated by the concentrated localization of ATN-1 (Fig. 1A). This pattern is consistent with previous reports (Ono *et al*., 1999; Ono *et al*., 2006; Yamashiro *et al*., 2007; Ono, 2024). When the muscle was contracted, the bands of F-actin were clearly separated in the middle of the I-bands, where the rows of dense bodies were located (Fig. 1B). Therefore, the localization patterns of F-actin and ATN-1 were nearly complementary, and their co-localization, if any, was minimal.

**Fig. 1.**
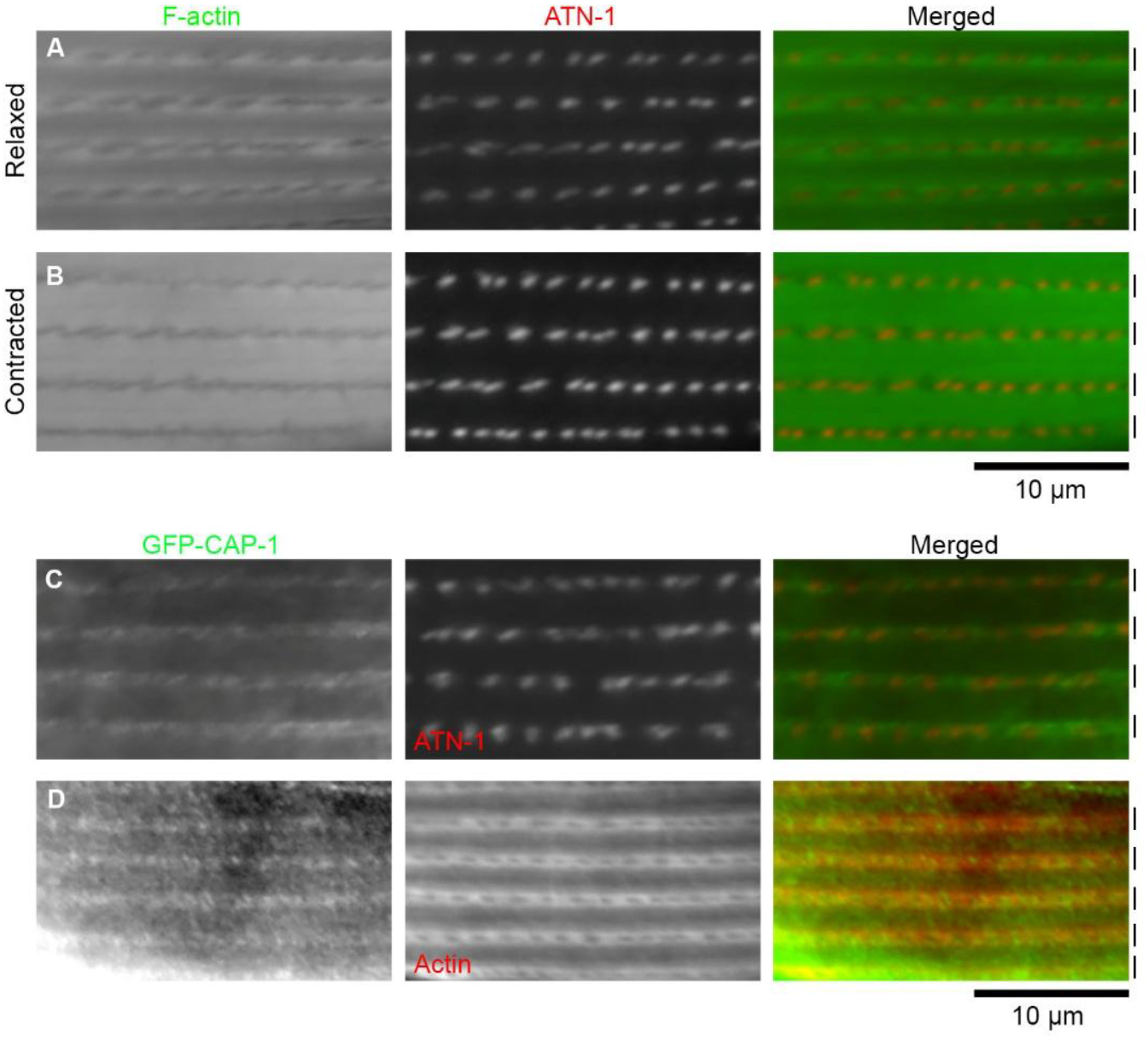
F-actin and CAP-1 are not concentrated at the ATN-1-enriched dense bodies. *A, B*, Localization patterns of F-actin (left) and ATN-1 (middle) were examined by staining with Alexa 488-labeled phalloidin and mCherry-tagging of ATN-1. Merged images are shown (right). Approximate positions of the I-bands are indicated by vertical bars on the right. With this procedure, both relaxed (*A*) and contracted (*B*) states of muscle cells were observed. Rows of the dense bodies are oriented horizontally. *C, D*, Localization of GFP-CAP-1 (left) was compared with that of ATN-1 (*C*, middle) or actin (*D*, middle) by immunostaining. Merged images are shown (right). With the immunostaining procedure, only relaxed states of muscle cells were observed. Bar, 10 µm.

### Capping protein is localized to linear arrays in an ATN-1-independent manner

To determine the precise location of the barbed ends of sarcomeric actin filaments, we examined localization of CP in the body wall muscle. In cross-striated muscle, CP is also known as CapZ, which means an actin capping protein at the Z-discs/lines (Casella *et al*., 1987). Also, CP exclusively binds to the barbed ends of actin filaments *in vitro* (Edwards *et al*., 2014). Therefore, CP is considered the best marker for the actin barbed ends. In *C. elegans*, α and β subunits of CP are encoded by *cap-1* and *cap-2*, respectively, and the CAP-1/CAP-2 heterodimer caps the barbed ends of actin filaments *in vitro* (Waddle *et al*., 1993). A recent study has shown that CAP-1 is expressed in the body wall muscle and localizes in a striated pattern (Ray *et al*., 2023), but precise localization of CAP-1 in sarcomeres has not been determined. We used a strain with an endogenous green fluorescent protein (GFP) tag on the *cap-1* gene (Yan *et al*., 2022) and confirmed that CAP-1 was expressed in the body wall muscle and localized in a striated pattern (Fig. 1C, D). Localization of CAP-1 was uniformly linear (Fig. 1C, D). By comparing with the ATN-1 localization, CAP-1 appeared partially overlapped with ATN-1, but their colocalization was minimal (Fig. 1C), indicating that CP-capped actin barbed ends are not converged on the α-actinin-rich dense bodies. By comparing with the actin localization, CAP-1 was concentrated in the middle of the I-bands (Fig. 1D). Unlike the phalloidin staining, immunofluorescent staining of actin appeared mostly in a relaxed chain-like pattern (Fig. 1D, middle), even when worms had been treated with levamisole to induce muscle contraction immediately before the fixation, suggesting that the contracted state was not preserved well in our immunostaining procedures (our unpublished observations). These localization patterns demonstrate that the CP-capped actin barbed ends are linearly aligned in the middle of the I-bands without directly converging on the dense bodies.

These microscopic data strongly suggest that the barbed ends of the sarcomeric actin filaments are not directly anchored to the dense bodies (Fig. 2). Rather, the CP-capped actin barbed ends are aligned in a linear Z-line-like pattern without converging on the dense bodies as illustrated in Fig. 2. When the muscle is relaxed, the stagger angle is shallow, and the actin filaments are laterally close to the Z-line-like structures (Fig. 2A), which may give an appearance of a chain-like pattern. When the muscle is contracted, the stagger angle is increased, and the actin filaments are pulled by myosin and move away from the Z-line-like structures (Fig. 2B), generating clear gaps of F-actin in the middle of the I-bands as seen in Fig. 1B.

**Fig. 2.**
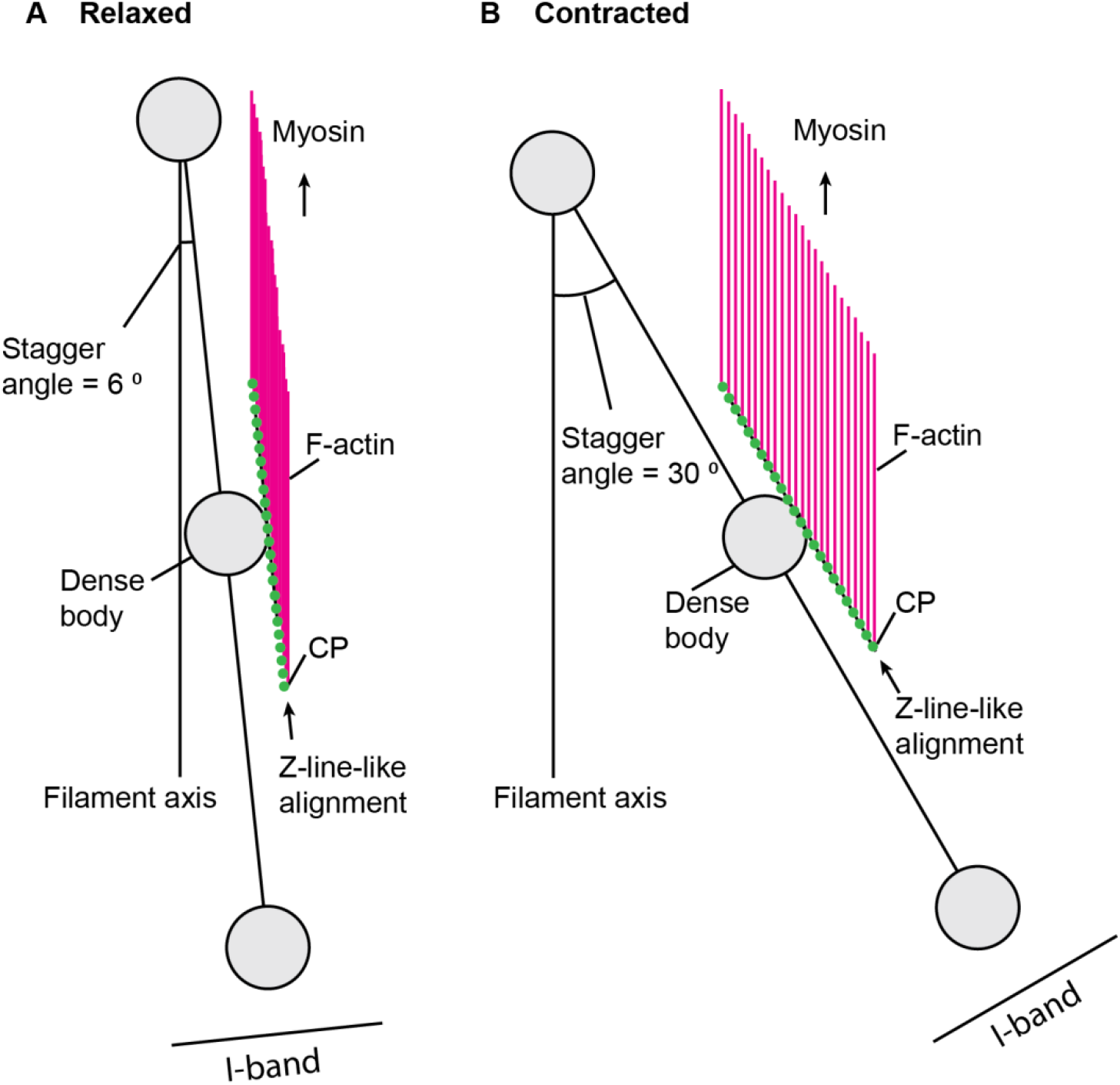
A model of organization of actin filaments and CP in the *C. elegans* body wall muscle. This model is a view from the top (the body surface). The axis of thin and thick filaments is oriented vertically. The oblique angle of the dense body alignment to the filament axis is termed “stagger angle”. The stagger angle is small at a relaxed state (6 ⁰ in *A*) but large at a contracted state (30 ⁰ in *B*). The scale of the illustration is based on the representative size of the dense bodies (0.2 µm) and distance between the nearest dense bodies (1.0 µm). CP (green dots) caps the barbed ends of F-actin (magenta) and is linearly arranged in a Z-line-like alignment. F-actin remains parallel at both states. For simplicity, only actin filaments in the top halves near the dense bodies in the center are shown.

The relationship between the linear arrangement of CAP-1 and the dense bodies were further examined using an α-actinin mutant. Knockout of the *atn-1*/α-actinin gene causes significant shortening of the cytoplasmic portion of the dense bodies and formation of actin aggregates with mild decrease in muscle contractility but does not cause major effects on sarcomere integrity (Moulder *et al*., 2010). In the *atn-1-null* mutant, CAP-1 still localized to a linear pattern, which was indistinguishable from the pattern in wild-type background (Fig. 3, low-magnification images in A and B, high-magnification images in C and D). CAP-1 was not concentrated in the actin aggregates in the *atn-1-null* mutant (Fig. 3B, arrows). Quantitative analyses indicated that actin aggregates were detected in almost all *atn-1* mutants but nearly none in wild-type (Fig. 3E), and that linear CAP-1 arrangement was detected in almost all wild-type and *atn-1* mutant animals (Fig. 3F). Therefore, the aggregated actin might be randomly polymerized without proper capping by CP. These results demonstrate that the linear arrangement of CAP-1 is maintained independently of ATN-1 and the major cytoplasmic portion of the dense bodies, which explains why *atn-1-null* mutation causes only mild sarcomere defects (Moulder *et al*., 2010).

**Fig. 3.**
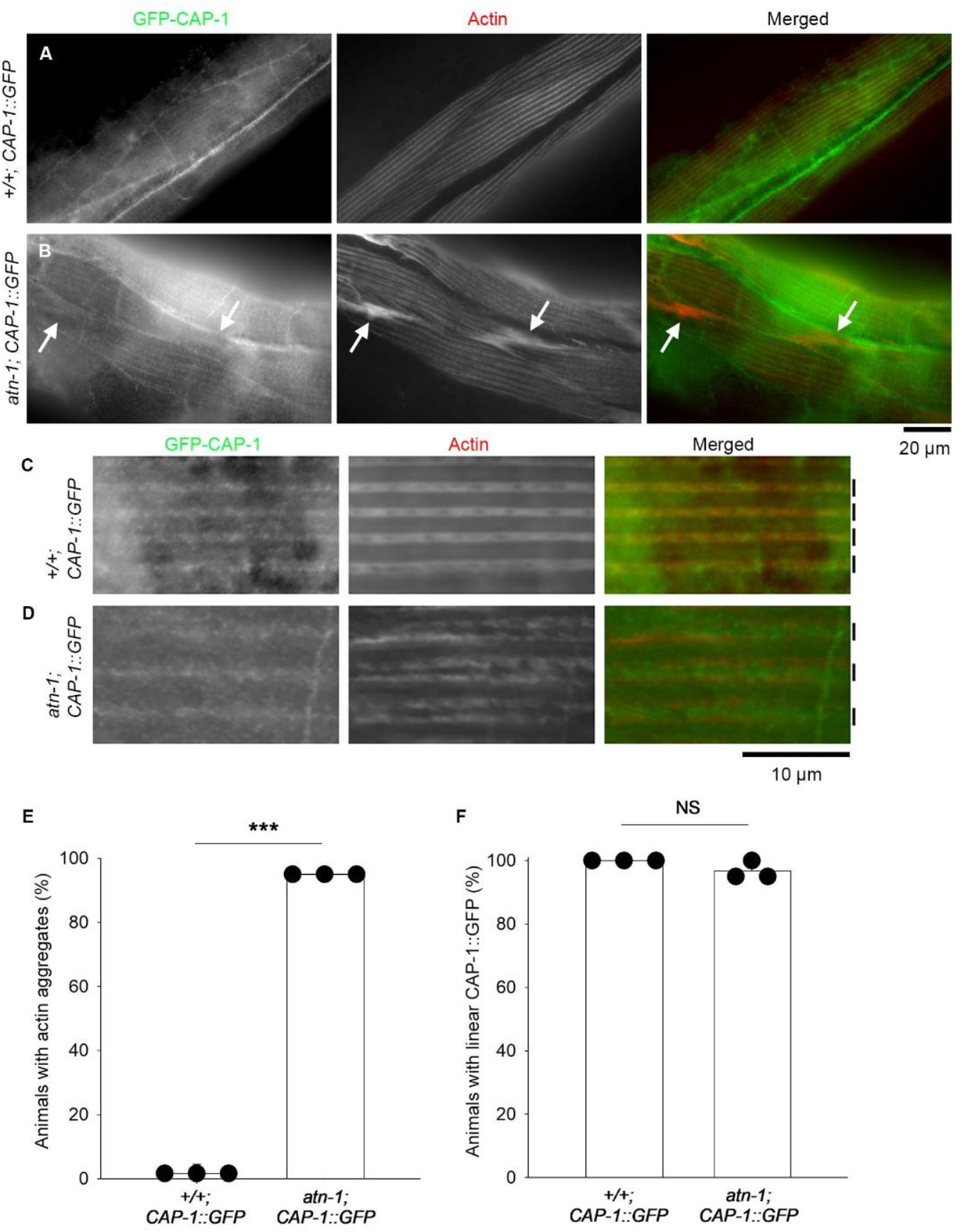
Linear alignment of CAP-1 is maintained independently of ATN-1. *A-D*, Localization patterns of GFP-CAP-1 (left) and actin (middle) were examined in wild-type (*+/+*) and *atn-1-null* (*atn-1*) backgrounds. Merged images are shown (right). In low-magnification views (*A, B*; Bar, 20 µm), linear striated pattern of GFP-CAP-1 (*A*) was not disrupted in *atn-1* (*B*). Actin aggregates were formed in *atn-1* (*B* middle, arrows), but GFP-CAP-1 was absent from the aggregates (*B* left, arrows). In high-magnification views (*C, D*; Bar, 10 µm), the linear alignment of GFP-CAP-1 was detected at the middle of the I-band (approximate positions indicated by vertical bars on the right) in both wild-type (*C*) and *atn-1* (*D*). *E, F*. Quantification of percentages of animals with actin aggregates (*E*) and with normal linear GFP-CAP-1 alignment (*F*). n = 3. ***, p < 0.001. NS, not significant.

### Capping protein is required for proper assembly of sarcomeric actin filaments and embryonic or larval development

Depletion of the CP subunits by RNA interference (RNAi) caused embryonic or larval lethality with severe sarcomere defects in the body wall muscle (Fig. 4 and 5), indicating that CP is essential for proper sarcomere assembly. When *cap-1* or *cap-2* was knocked down by RNAi, ∼70 % of the animals were arrested at embryonic or L1 larval stage (Fig. 4B-D) when control worms were fully grown to adults (Fig. 4A). Both *cap-1(RNAi)* and *cap-2(RNAi)* produced indistinguishable phenotypes (Fig. 4B-D). To determine the muscle phenotypes in embryos and larvae, the RNAi-treated animals were examined by staining with fluorescently labeled phalloidin to visualize F-actin organization (Fig. 5). In control embryos, at the two-fold stage (∼450-min-old embryos), various sizes of F-actin bundles were formed in a somewhat disorganized manner (Fig. 5A, top). Then, at the three-fold stage (∼520-min-old embryos), uniform F-actin bundles were linearly assembled into sarcomeres (Fig. 5 B and C, top). These patterns of F-actin are consistent with the previous reports of muscle actin assembly in wild-type embryos (Epstein *et al*., 1993; Hresko *et al*., 1994; Ono *et al*., 1999). In both *cap-1(RNAi)* and *cap-2(RNAi)* embryos, the appearance of various sizes of F-actin bundles was not clearly different from those in control embryos at the two-fold stage (Fig. 5A, middle and bottom). However, F-actin remained disorganized and failed to be assembled in a linear uniform pattern at the three-fold stage (Fig. 5B and C, middle and bottom). Similar phenotypes persisted in the L1 larval stage (Fig. 5D). Disorganized F-actin in the body wall muscle was detected in ∼70 % of *cap-1(RNAi)* and *cap-2(RNAi)* animals but rarely found in control animals (Fig. 5E). These results indicate that CP is required for organized uniform assembly of sarcomeric actin filaments in a late embryonic stage but not essential for initial assembly of actin filaments. Vertebrate CPs have been demonstrated to be important for sarcomeric actin assembly in skeletal muscles (Schafer *et al*., 1995; Berger *et al*., 2022). Accordingly, the essential role of CP in sarcomere assembly is conserved in the obliquely striated muscle of the nematode *C. elegans*.

**Fig. 4.**
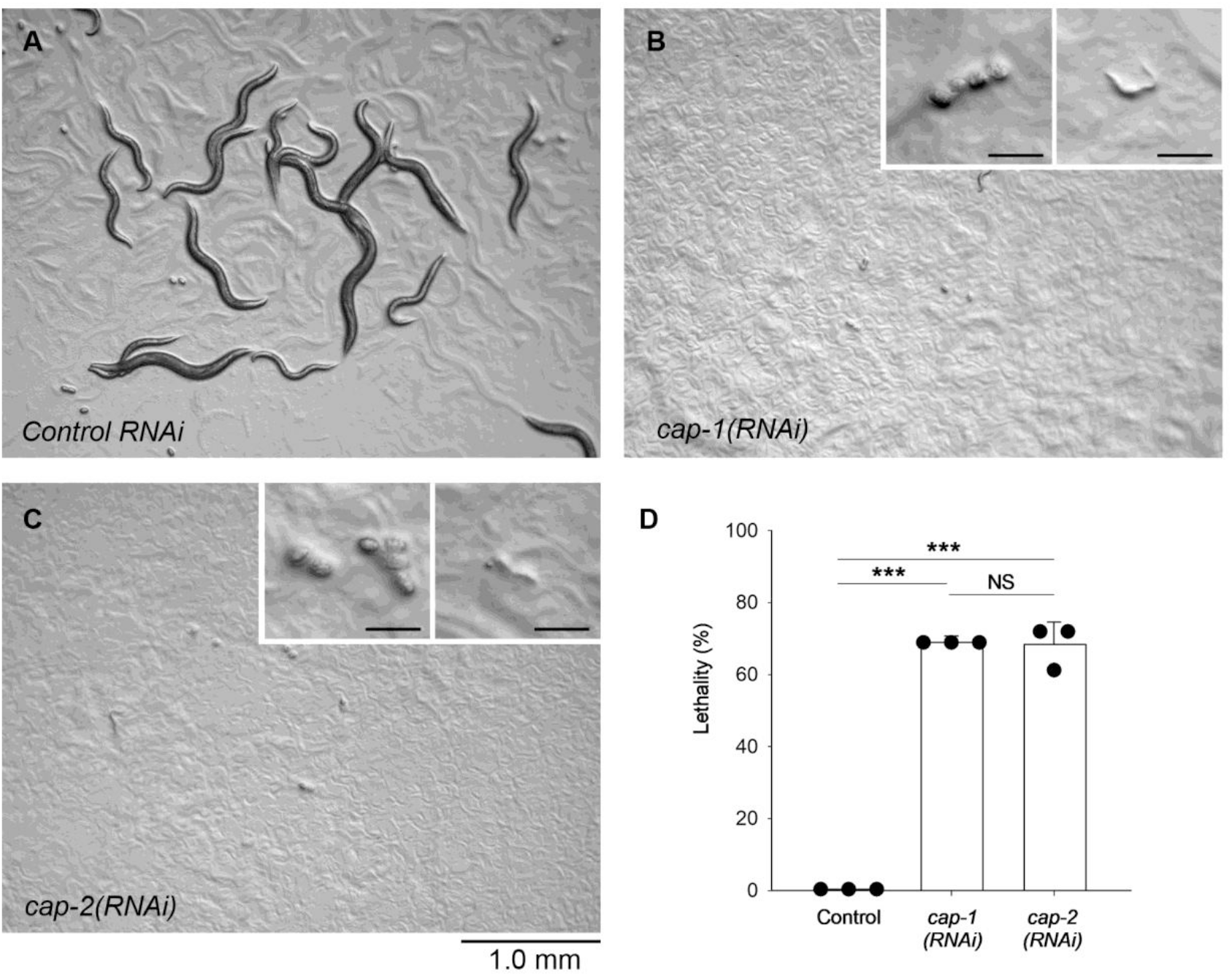
RNA interference of genes encoding CP subunits, *cap-1* and *cap-2*, causes embryonic or larval lethality. Worm cultures treated with control RNAi (*A*), *cap-1(RNAi)* (*B*), or *cap-2(RNAi)* (*C*) were observed on agar plates. Bar, 1.0 mm. At a time when most of control worms grew to adults (*A*), most of *cap-1(RNAi)* (*B*) and *cap-2(RNAi)* (*C*) animals were arrested at embryos or early larvae (insets in *B* and *C*, Bars, 0.1 mm). *D*, Lethality was quantified after 3 days of RNAi treatments when most of control worms reached the adult stage. Animals that were arrested at embryonic or early larval stages were counted as dead. n = 3. ***, p < 0.001. NS, not significant.

**Fig. 5.**
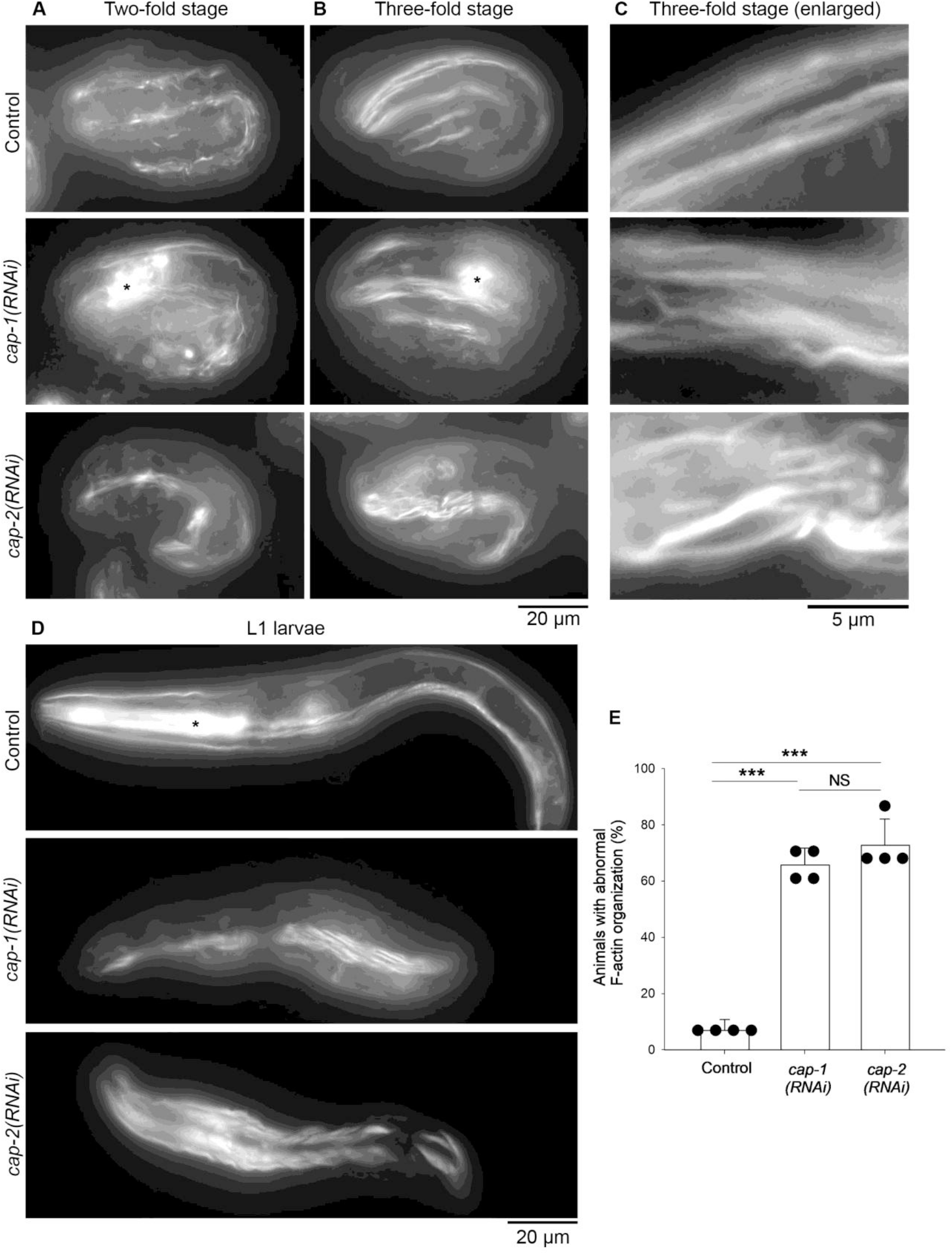
RNA interference of *cap-1* or *cap-2* causes disorganization of actin filaments in embryonic and larval body wall muscle. Worm cultures treated with control RNAi, *cap-1(RNAi)*, or *cap-2(RNAi)* were stained with rhodamine-phalloidin. *A-D*, Representative images of two-fold embryos (∼450-min-old) (*A*), three-fold embryos (∼520-min-old) (*B*, enlarged images shown in *C*), and L1 larvae (*D*) are shown. The heads are oriented to the left. Asterisks indicate fluorescence from the pharyngeal muscle. *E*, Quantification of percentages of animals with abnormal F-actin organization. n = 4. ***, p < 0.001. NS, not significant. Bars, 20 µm (*A, B*, and *D*) and 5 µm (*C*)

### Perspectives on the relationship between the dense bodies and the Z-line-like structures in obliquely striated muscle

In contradiction to the previous assumption, our results indicate that the thin filaments of the *C. elegans* body wall muscle are not directly anchored to the α-actinin-rich dense bodies. Therefore, we propose that the dense bodies of the *C. elegans* body wall muscle are not equivalent to the Z-discs/lines of transversely cross-striated muscles (Fig. 2). However, actomyosin-generated contractile forces still need to be transmitted to the muscle cell membrane and further external hypodermis and cuticles for body movement (Burr and Gans, 1998). The base of the dense bodies contains integrin-based attachments and considered equivalent to the costameres of vertebrate cross-striated muscles (Ervasti, 2003; Cox and Hardin, 2004; Lecroisey *et al*., 2007; Gieseler *et al*., 2017). Actomyosin contraction causes deformation of the cytoplasmic portions of the dense bodies (Kimmich *et al*., 2024), indicating that contractile forces are applied to the dense bodies. Therefore, actin filaments should be physically linked indirectly to the cytoplasmic portions of the dense bodies by an unknown mechanism. Intriguingly, some proteins specifically localize to gaps between dense bodies in the *C. elegans* body wall muscle, such as UNC-60B (actin depolymerizing factor/cofilin) (Ono *et al*., 1999), KETN-1 (kettin) (Ono *et al*., 2006; Ono *et al*., 2020), CAS-1 (cyclase-associated protein) (Nomura *et al*., 2012), FHOD-1 (formin) (Mi-Mi *et al*., 2012), LIM-8 (Qadota *et al*., 2007), and DBN-1 (actin-binding protein-1) (Butkevich *et al*., 2015). Most of these proteins directly bind to actin, and their locations overlap with that of CP. Therefore, these are candidate components of the mechanism to link sarcomeric actin filaments to the dense bodies, but further investigation is needed to understand the molecular mechanism of such linkages.

It is surprising that *C. elegans* ATN-1/α-actinin in the dense bodies is largely not engaged directly with F-actin or CP, since vertebrate α-actinin is the major F-actin-bundling protein in the Z-discs/lines of vertebrate cross-striated muscles (Masaki *et al*., 1967) and binds directly to CP (Papa *et al*., 1999). *C. elegans* ATN-1 may have functions other than F-actin bundling. Human α-actinin-2 (a major muscle isoform) binds to various cytoskeletal and non-cytoskeletal proteins (Ladha *et al*., 2021) and form phase-separated condensates with FATZ-1 (Sponga *et al*., 2021). Similarly. *C. elegans* ATN-1 may contribute to generating compartmentalized environments in the form of the dense bodies to concentrate certain proteins other than actin. Thus, additional biochemical and genetic studies on ATN-1 are required to understand whether ATN-1 has currently unknown functions. In conclusion, we propose to revise the previous notion that ATN-1 directly bundles actin filaments at the dense bodies and to initiate investigation on the molecular mechanism of the linkage between actin and the dense bodies as well as novel actin-independent function of ATN-1.

## Materials and methods

### *C. elegans* strains and culture

The worms were cultured following standard methods (Stiernagle, 2006). Wild-type N2 and RB1812 *atn-1(ok84)* (Moulder et al., 2010) were obtained from the *Caenorhabditis* Genetics Center (Minneapolis, MN). RSL62 *atn-1(ftw35 [atn-1::mCH::ICR::GFPnls])* (Kimmich et al., 2024) was provided by Drs. Ryan Littlefield (Ohio University, Athens, OH) and David Pruyne (State University of New York, Syracuse, NY). SWG061 *cap-1(ges4[cap-1::GFP + LoxP]); gesIs003[Pmex-5::Lifeact::mKate2::nmy-2UTR,unc-119+]* (Yan et al., 2022) was provided by Dr. Stephan W. Grill (Max Planck Institute of Molecular Cell Biology and Genetics, Dresden, Germany). ON398 *cap-1(ges4[cap-1::GFP + LoxP]); atn-1(ok84)* was generated by crossing SWG061 and RB1812 and isolating homozygous progeny.

### Fluorescence microscopy

Staining of whole worms with Alexa 488–phalloidin (Catalog number A12379, Thermo Fisher Scientific) or tetramethylrhodamine-phalloidin (Catalog number P1951, MilliporeSigma) was performed as described previously (Ono, 2001, 2022). Immunofluorescent staining of whole worms was performed as described previously (Nonet *et al*., 1997). Primary antibodies used were mouse anti-actin monoclonal (C4, Catalog number MA5-11869, Thermo Fisher Scientific), rabbit anti-GFP polyclonal (Catalog number A-11122, Thermo Fisher Scientific), and mouse anti-ATN-1 monoclonal (MH35, provided by Dr. Pamela Hoppe, West Michigan University) (Francis and Waterston, 1985). Secondary antibodies used were Alexa 488-labeled goat anti-rabbit IgG (Catalog number A-11008, Thermo Fisher Scientific) and Cy3-labeled donkey anti-mouse IgG (Catalog number 715-165-151, Jackson ImmnoResearch).

Samples were mounted with ProLong Diamond Antifade Mountant (Catalog number P36970, Thermo Fisher Scientific) and observed by epifluorescence using a Nikon Eclipse TE2000 inverted microscope (Nikon Instruments, Tokyo, Japan) with a CFI Plan Apo Lambda 100x (oil, NA 1.45) objective. Images were captured by a Hamamatsu ORCA Flash 4.0 LT sCMOS camera (Hamamatsu Photonics, Shizuoka, Japan) and processed by NIS-Elements (Nikon Instruments) and Adobe Photoshop 2026.

### RNA interference

RNAi experiments were performed by feeding with *Escherichia coli* HT115 expressing double-stranded RNA as described previously (Ono and Ono, 2002). An RNAi clone for *cap-1* (IV-3J07) (Kamath *et al*., 2003) was obtained from MRC Geneservice (Cambridge, United Kingdom). An RNAi clone for *cap-2* (mv_CAA79270) (Rual *et al*., 2004) was obtained from Dharmacon/Thermo Fisher Scientific. An empty vector L4440 (provided by Dr. Andrew Fire, Stanford University) was used as a control. L4 larvae were treated with RNAi, and phenotypes were characterized in the F1 generation.

### Statistics

Data were analyzed by Student’s *t*-test (Fig. 3E, F) or one-way analysis of variance with the Holm–Sidak method for pairwise comparison (Fig. 4D; Fig. 5E) using SigmaPlot 15.0 (Grafiti LLC).

## Abbreviations

CP: capping protein
GFP: green fluorescent protein
RNAi: RNA interference

## Acknowledgements

We thank Dr. Guy Benian for valuable comments on the manuscript. Some *C. elegans* strains were provided by the *Caenorhabditis* Genetics Center, which is funded by the National Institutes of Health Office of Research Infrastructure Programs (P40 OD010440). This work was supported by a grant from the National Institutes of Health (R01-GM144563) to S. O.

